# A defined community of core gut microbiota members promotes cognitive performance in honey bees

**DOI:** 10.1101/2023.01.03.522593

**Authors:** Amélie Cabirol, Julie Schafer, Nicolas Neuschwander, Lucie Kesner, Joanito Liberti, Philipp Engel

## Abstract

The composition of the gut microbiota has recently been identified as a cause of cognitive variability in humans and animals. Germ-free individuals and individuals exposed to an antibiotic treatment show severe alteration of their learning and memory performance measured in various cognitive tasks. While different species of bacteria are known to interact in the gut, their cumulative or synergistic effects on cognitive performance remain elusive. Here we established a defined bacterial community - composed of core members of the corbiculate bee microbiota - which enhances honey bees’ cognitive capacities. Honey bees colonized with this reconstituted community discriminated better two odours based on the presence or absence of a sucrose reward compared to germ-free individuals. They also memorized better these odour-food associations in the short-term. These cognitive improvements seem to constitute an emergent property of the community, because they could not be recapitulated by any of the community members when mono-associated in gnotobiotic bees and they were not explained by the total biomass in the gut. The identification of this community and its effect on bees open new avenues of research in neuroscience, microbiology, and ecology. Future research should help understanding how interactions between bacterial species in the community promote the host’s associative learning and memory performance.

## Introduction

Evidence for a critical role of the gut microbiota in modulating cognitive performance has been accumulating over the past decade in humans and laboratory animals^1,2^. Different gut microbiota profiles have been associated with different learning scores in 2-year-old human infants, suggesting a role of the gut microbiota in cognitive development^3^. In adults, obesity-associated cognitive deficits correlated with increased relative abundance of actinobacteria in the gut^4^. Beyond associations, the use of reductionist laboratory animals have allowed inferring a causal link between a healthy gut microbiota and normal cognitive performance^5,6^. Germ-free (GF) mice and mice treated with antibiotics exhibited deficits in associative memory and extinction learning following a pavlovian cued fear conditioning where a sound was associated with an electric shock^7,8^. While studies in GF animals have revealed a profound impact of gut microbiota deprivation on cognitive performance, the contribution of specific gut members and microbial communities to cognitive functions has mostly remained elusive^2,5^. An attempt to restore fear extinction learning deficits in ex-GF mice using a simplified community of gut symbionts has failed^8^ and further investigations are therefore needed to understand the mechanisms by which gut bacteria affect cognitive functions.

The honey bee (*Apis mellifera*) is emerging as a useful model to unravel the proximate mechanisms underlying the effect of the microbiota on complex cognitive and behavioural phenotypes^9–11^. Despite a relatively simple brain (1 million neurons), honey bees exhibit impressive cognitive skills that have been studied under laboratory conditions for over 60 years^12,13^. Honey bee workers harbour a simple and natural gut community of 8-10 bacterial phylotypes which can be cultured under laboratory conditions and inoculated in GF bees^14,15^. Five of these phylotypes (Gilliamella, Snodgrassella, Bombilactobacillus Firm-4, Lactobacillus Firm-5, and Bifidobacterium) constitute the corbiculate core microbiota, as they are present in almost every adult female worker bee of *A. mellifera* and are also found in other corbiculate bees including other honey bee species, bumblebees and stingless bees^16^. A recent study has shown that GF bees display altered associative olfactory memory capacities^10^. Mono-colonization with the bacteria *Lactobacillus* Firm-5 did not improve bees’ memory retention of the odour-food association unless bees were supplemented with tryptophan. Although these results confirm those observed in GF laboratory rodents and identify a possible mechanism, the relative contribution of different gut bacteria or communities of bacteria to cognitive improvements is still unknown.

Here we assessed the contribution of the corbiculate core microbiota to honey bees’ associative learning and memory abilities using a pavlovian olfactory conditioning assay. We successfully generated gnotobiotic bees, whose gut was either GF, colonized by a reconstituted community of 11 strains representing the five core members of the honey bee gut microbiota (BeeCom_001) or mono-colonized by each core member. Our results show that colonisation with the BeeCom_001 significantly improved bees’ olfactory learning and memory performance while none of the community members alone could recapitulate those effects.

## Results

### Quantitative PCR confirms the gnotobiotic status of the experimental bees

Gnotobiotic bees were obtained from 14 experimental replicates as previously described^17^ by feeding newly emerged GF individuals an inoculum of bacteria of interest (5 μl of OD_600_ = 0.1) and leaving them in cages for eight days. Each experiment included seven different treatment groups, i.e. GF bees, bees colonized with the defined community BeeCom_001, or bees colonized with each of the five core members separately: Gilliamella (Gi), Snodgrassella (Sn), Bombilactobacillus Firm-4 (F4), Lactobacillus Firm-5 (F5), or Bifidobacterium (Bi). The composition of the inoculum did not affect the sucrose consumption per bee and the mortality in the cages (Supplementary Figure 1). The guts of 8-day-old bees were dissected immediately after the olfactory conditioning assay, and the quantity of all bacteria and of each bacterial phylotype was assessed by quantitative PCR (qPCR) using universal 16S rRNA gene primers and phylotype-specific primers, respectively. The weight of individual bees (without gut) was not different between the treatment groups (Supplementary Figure 1).

The qPCR analyses validated the gnotobiotic status of our bees and confirmed that the different bacterial treatments had worked (Figure 1). First, the total number of bacteria in the gut of GF bees (as assessed with the universal 16S rRNA gene primers) was significantly lower than of bees colonized with the BeeCom_001 or mono-colonized with individual core members (pairwise Wilcoxon Rank Sum tests corrected for the False Discovery Rate – FDR; p < 0.0001 for all comparisons). Second, all mono-colonized bees had high levels of the treatment-specific community member in their gut (as assessed with the phylotype-specific primers), and in GF bees, their levels were mostly below the detection limit. And third, most bees of the BeeCom_001 treatment were abundantly colonized by each of the five different community members, with the exceptions of a few bees for which no or very low levels of Gilliamella, Snodgrassella, or Bifidobacteria were detected. Notably, bees mono-colonized with Gilliamella, Snodgrassella or Bifidobacterium had significantly lower total bacterial loads than bees colonized with the BeeCom_001 (BeeCom_001 vs. Gi: p < 0.005; BeeCom_001 vs. Sn: p < 0.01; BeeCom_001 vs. Bi: p < 0.05), whereas the total bacterial levels of bees mono-colonized with Bombilactobacillus Firm-4 or Lactobacillus Firm-5 were similar to those colonized with the BeeCom_001.

**Figure 1.**
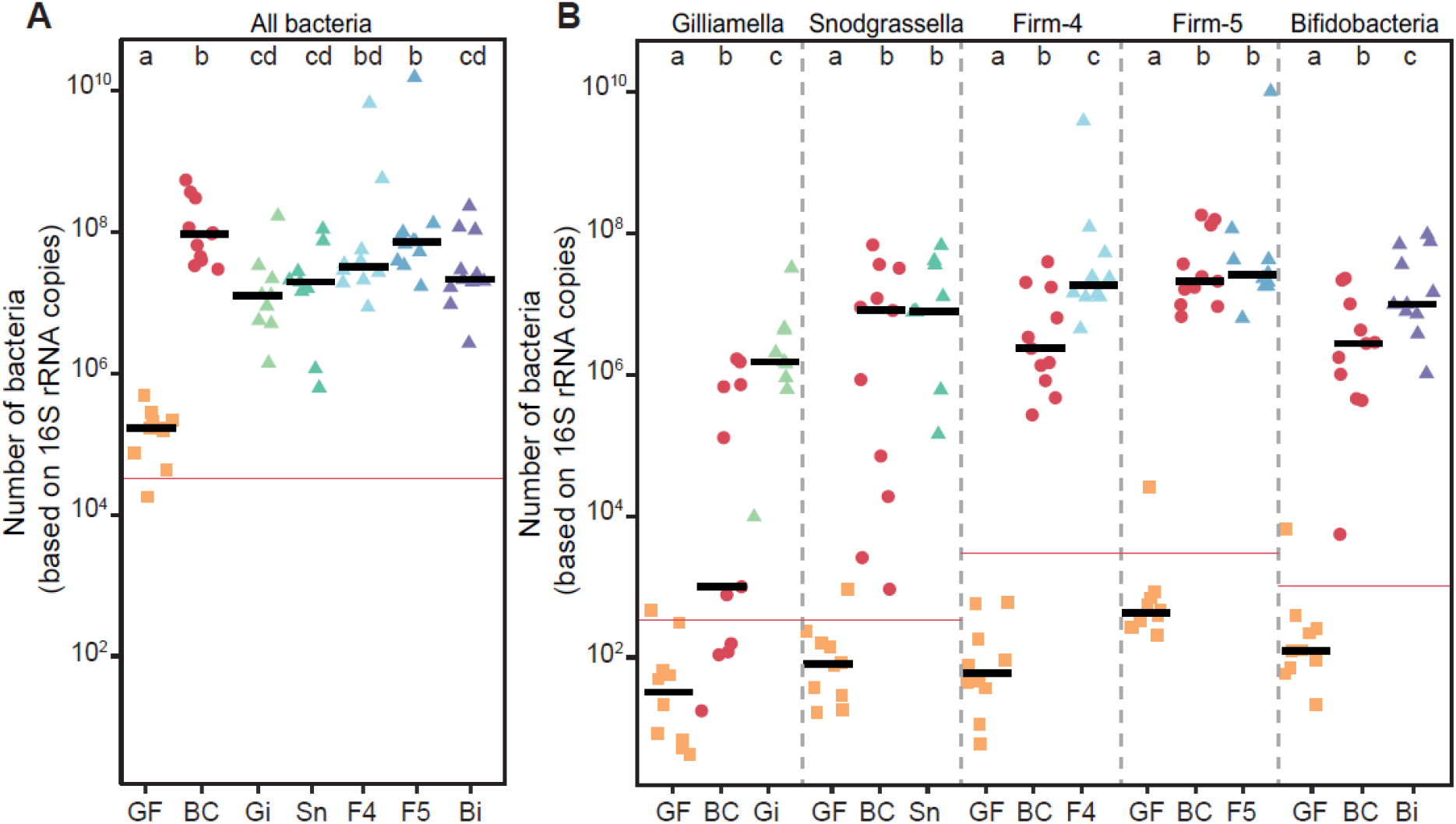
Bacterial loads in the gut of germ-free bees (GF; ■), bees colonized with the defined community BeeCom_001 (BC; ●) composed of the five core members of the bee gut microbiota, and bees mono-colonized (▲) with one of the five core members: Gilliamella (Gi), Snodgrassella (Sn), Bombilactobacillus Firm-4 (F4), Lactobacillus Firm-5 (F5), or Bifidobacterium (Bi). The number of bacteria was measured by qPCR using universal 16S rRNA gene primers (all bacteria) or species-specific primers for the same bee. The red lines represent the limit of detection of each primer pair under which bacterial loads were too low to be accurately measured. Different letters indicate significant differences between gut conditions for each primer pair (Wilcoxon Rank Sum test, false-discovery rate correction).

### Learning and memory was improved by the defined core community of gut bacteria

Olfactory learning and memory performance was assessed in 8-day-old bees using the well-established classical differential conditioning of the proboscis extension response (PER; Figure 2A)^18,19^. In this assay, the unconditioned stimulus (US) consisted in sucrose solution that, when applied to a bee’s antennae, automatically triggered the PER. While one odour was positively reinforced by being presented in association with the US (conditioned stimulus, CS+), a second odour was not reinforced (CS-). Proboscis extension responses to the CS were recorded as a measure of learning (conditioned responses). Bees received five presentations of each CS in a randomized order. Short-term memory was assessed 15 min after the end of conditioning by presenting both CS in absence of any US.

**Figure 2.**
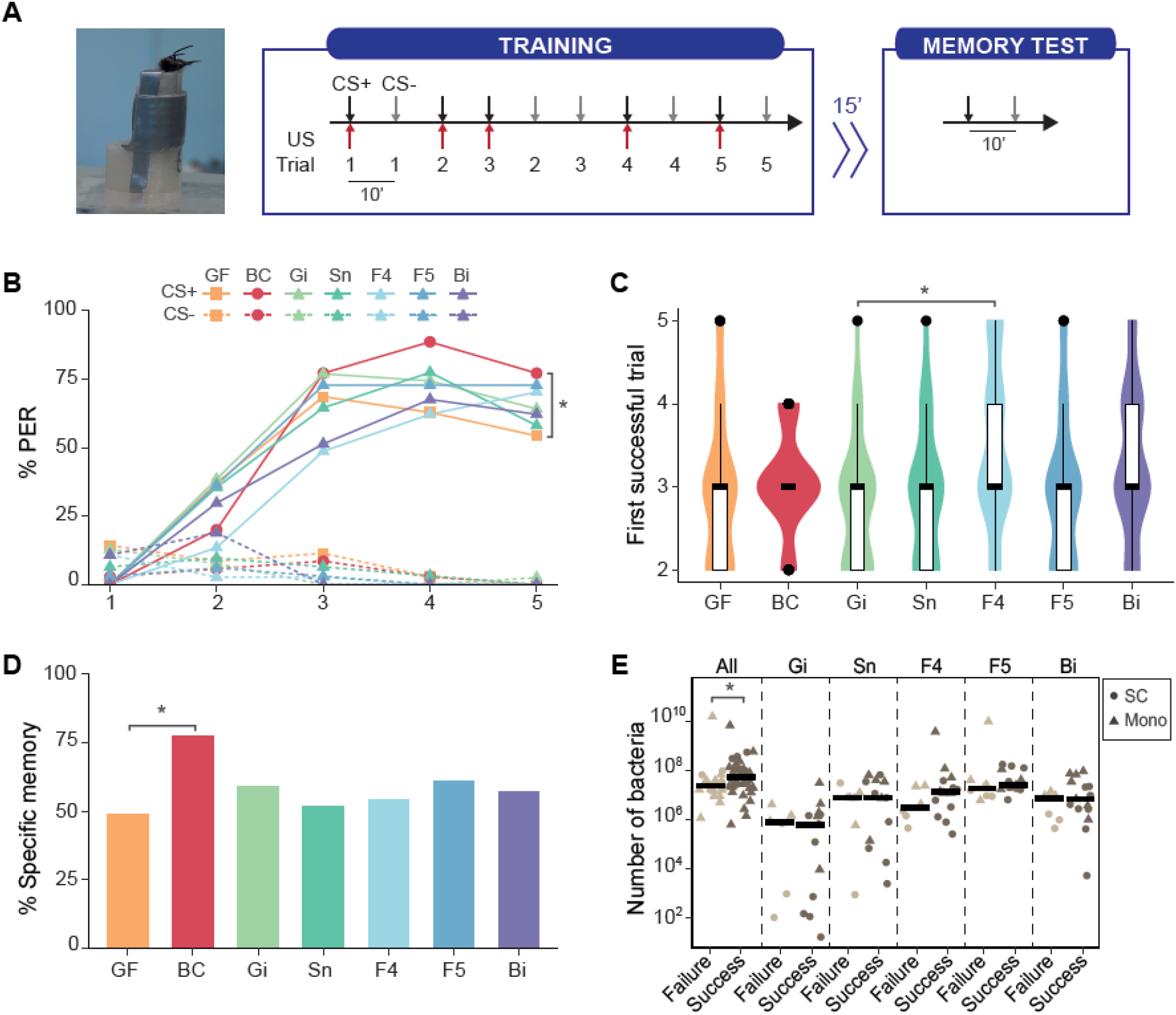
The defined core community of gut bacteria facilitated olfactory learning and memory performance of their honey bee host. **A.** Olfactory differential conditioning of the PER protocol. The picture shows a harnessed honey bee extending its proboscis. Bees were trained to discriminate an odour (conditioned stimulus, CS+) rewarded with sucrose solution (unconditioned stimulus, US) from an unrewarded odour (CS-) within 5 presentations of each. Short-term memory was assessed 15 minutes after the end of the conditioning by presenting both CS in absence of the US. **B.** Differential learning performance of germ-free bees (GF; n = 35), bees colonized with the defined community BeeCom_001 (BC; n = 35), and bees mono-colonized with Gilliamella (Gi; n = 39), Snodgrassella (Sn; n = 31), Bombilactobacillus Firm-4 (F4; n = 37), Lactobacillus Firm-5 (F5; n = 33), or Bifidobacterium (Bi; n = 37). The percentage of bees exhibiting PER to the CS+ (plain lines) and CS-(dashed lines) is displayed across the 5 learning trials. **C.** First successful trial of gnotobiotic bees. **D.** Percentage of gnotobiotic bees responding to the CS+ but not to the CS-during the memory test. **E.** Number of bacteria in the gut of non-GF bees (● = BC; ▲ = mono-colonized) who succeeded or failed the memory test. The number of bacteria was measured by qPCR using universal 16S rRNA primers (all bacteria) and phylotype-specific primers. (*) p < 0.05.

All groups of gnotobiotic bees solved the differential learning task but exhibited different learning patterns (Figure 2B-C). The percentage of bees showing PER to the CS+ increased across the 5 learning trials and differed significantly from the CS- (Figure 2B; Generalized Linear Mixed Model, GLMM; *CS × Trial*: p < 0.0001 for all gut conditions). The gut condition significantly influenced responses to the CS across trials (GLMM; *Gut × Trial*: χ^2^ = 13.39, df = 6, p < 0.05). In the last trial, the percentage of bees responding to the CS+ was significantly higher when the gut was colonized with the BeeCom_001 compared to when it was GF (Tukey’s post hoc tests, p < 0.05). None of the other gut conditions, at any trial, differed in their responses to the CS+ (p > 0.05 for all pairwise comparisons). Responses to the CS-were not affected by the gut condition (p > 0.05 for all pairwise comparisons). The speed of learning, reflected by the first trial at which a bee responded to the CS+ but not to the CS-, was also significantly influenced by the gut microbiota composition (Figure 2C; Kruskal-Wallis: χ^2^ = 16.84, df = 6, p < 0.01). Bees mono-colonized with Bombilactobacillus Firm-4 required more trials to solve the task compared to bees mono-colonized with Gilliamella (Dunn test corrected with the FDR; p < 0.05) and a similar trend was observed when compared to GF bees (p = 0.053), and to bees mono-colonized with Snodgrassella (p = 0.053) or Lactobacillus Firm-5 (p = 0.053). The first successful trial of bees colonized with the BeeCom_001 did not differ significantly from either GF bees (p = 0.43), or any mono-colonized condition (p > 0.05), suggesting an intermediate learning speed.

Short-term specific memory was improved by the gut colonization with the BeeCom_001 (Figure 2D). The percentage of bees colonized with the BeeCom_001 discriminating the CS+ from the CS-in the memory test was significantly higher than that of GF bees (GLM: estimates ± standard error = 1.27 ± 0.53, p < 0.05). None of the other gut conditions influenced memory performance (Tukey’s post hoc tests, p > 0.05). To assess whether the total biomass can explain differences in memory performance independently of the bacterial treatment, we compared the number of bacteria in the gut of non-GF bees according to their success or failure in the memory test (Figure 2E). The total number of bacteria (as assessed with universal 16S rRNA gene primers) in the gut of bees who successfully solved the memory test was larger than the one found in the gut of unsuccessful individuals (Wilcoxon Rank Sum tests, p < 0.05). None of the single phylotype abundances predicted memory performance (p > 0.05 for all phylotypes).

### Gut microbiota composition did not affect sucrose responsiveness

Sucrose responsiveness was shown to positively correlate with associative learning performance in conditioning protocols involving a sucrose reward as the US^20^. We therefore measured sucrose responsiveness to discriminate effects of gut bacteria on the perception of the US from effects on learning and memory *per se*. Following a standard protocol, proboscis extension was recorded in response to an ascending concentration series of six sucrose solutions, interspersed with water stimulations applied to the bees’ antennae^20^ (Figure 3). Responses to water can reflect the overall arousal or thirst of the tested individuals, as well as the phenomenon of sensitization occurring when individuals are repeatedly stimulated. This assay was performed prior to the olfactory conditioning.

**Figure 3.**
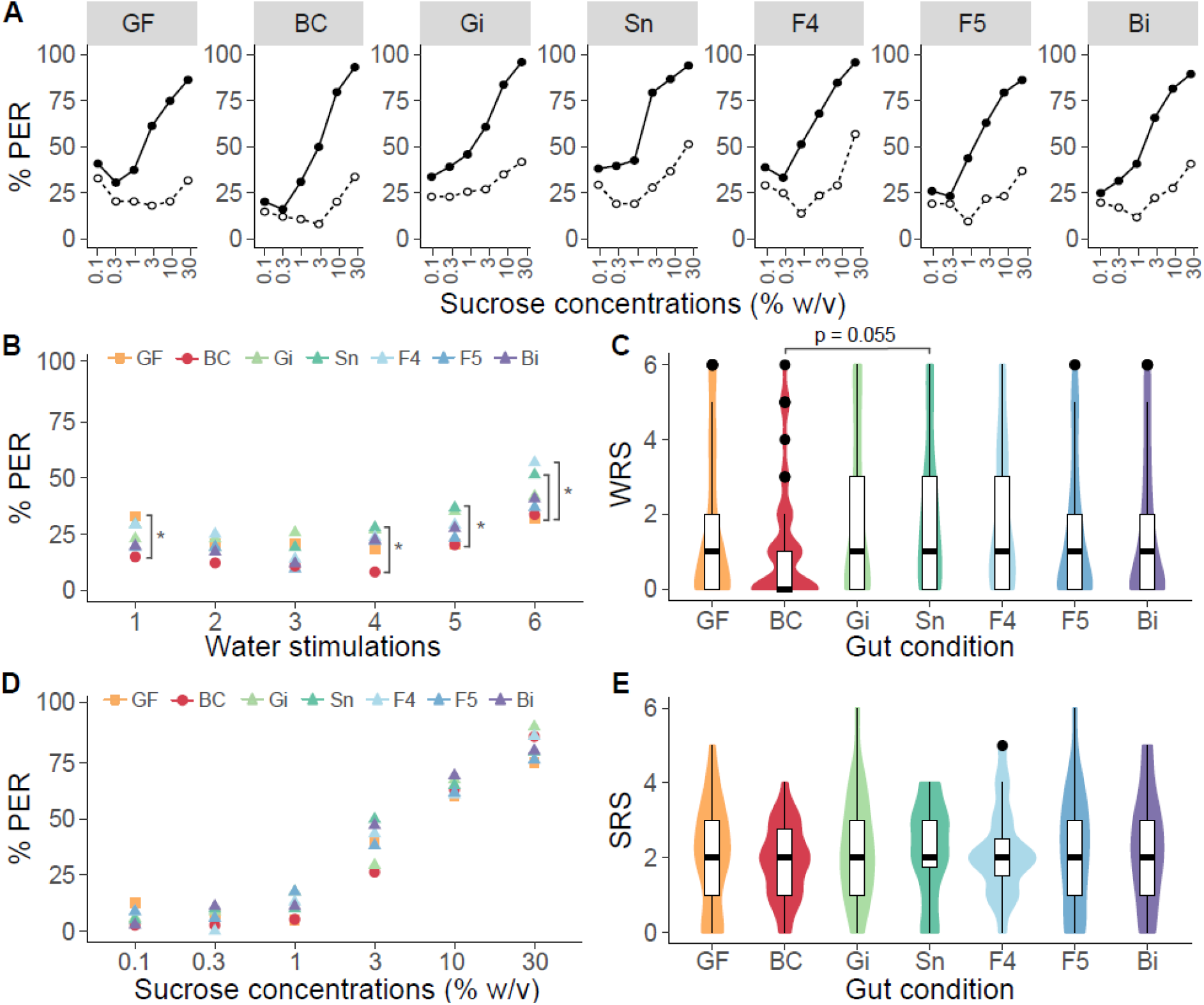
Gut bacteria influenced the water but not the sucrose responsiveness of their honey bee host. **A.** Percentage of bees showing a proboscis extension response (PER) to an ascending concentration series of 6 sucrose solutions (●) interspersed with water stimulations (○) presented to their antennae. The gut conditions were germ-free (GF, n = 88), colonized with the defined community BeeCom_001 (BC, n = 74), and mono-colonized with Gilliamella (Gi, n = 74), Snodgrassella (Sn, n = 68), Bombilactobacillus Firm-4 (F4, n = 72), Lactobacillus Firm-5 (F5, n = 73), or Bifidobacterium (Bi; n = 76). **B.** Comparison of the percentage of bees showing PER to the water stimulations between gut conditions. **C.** Water response scores (WRS) of bees from the different gut conditions. **D.** Comparison of the percentage of PER to the sucrose stimulations between gut conditions for bees that did not respond to any water stimulation (GF, n = 40; BC, n = 38; Gi, n = 34; Sn, n = 20; F4, n = 23; F5, n = 34; Bi, n = 36). **E.** Sucrose response scores (SRS) of bees from the different gut conditions. (*) p < 0.05.

For all gut conditions, the percentage of bees responding to the sucrose stimuli increased across the 6 trials and differed significantly from the percentage of bees responding to the water stimuli, thereby showing that bees could overall discriminate the two stimuli (Figure 3A; GLMM, *Trial*: p < 0.0001, *Stimulus × Trial*: p < 0.0001 for all gut conditions). Yet, the gut condition influenced bees’ sensitivity to water (Figure 3B-C). The percentage of bees responding to water across the 6 stimulations was significantly affected by the gut condition (Figure 3B; GLMM, *Gut × Trial*: χ^2^ = 21.29, df = 6, p < 0.005). Germ-free bees responded significantly more to the first presentation of water compared to bees colonized with the BeeCom_001 (Tukey’s post hoc test, p < 0.05). Bees mono-colonized with Snodgrassella or Bombilactobacillus Firm-4 showed a stronger sensitization to the repeated water stimulations compared to bees colonized with the BeeCom_001 or GF bees as demonstrated by a higher percentage of PER to the last few stimuli (Tukey’s post hoc test, *Trial 4 and Trial 5*: Sn vs. BC: p < 0.05, *Trial 6*: Sn vs. GF and F4 vs. GF: p < 0.05). Consistently, the water response scores (WRS), corresponding to the number of water stimuli each bee responded to, were significantly affected by the gut condition (Figure 3C; Kruskal-Wallis: χ^2^ = 13.46, df = 6, p < 0.05). Gut colonisation with Sn tended to increase the WRS compared to the BeeCom_001 (Dunn’s post hoc test corrected with FDR: p = 0.055). To exclude any effect of sensitization, sucrose responsiveness was assessed on bees that did not respond to the water stimuli (Figure 3D-E). The percentage of bees responding to sucrose increased with ascending concentrations but did not differ between gut conditions (Figure 3D; GLMM, *Trial*: χ^2^ = 226.86, df = 1, p < 0.0001, *Gut × Trial*: χ^2^ = 9.25, df = 6, p = 0.16). As a result, the sucrose response scores (SRS) were not affected either by the gut condition (Figure 3E; Kruskal-Wallis: χ^2^ = 1.85, df = 6, p = 0.93).

Only a subset of bees could be tested in the olfactory conditioning assay due to the defined timings of the experimental protocol. The SRS, but not the WRS, differed significantly between gut conditions in this subset (Supplementary Figure S2; Kruskal-Wallis; SRS: χ^2^ = 16.53, df = 6, p < 0.05; WRS: χ^2^ = 9.04, df = 6, p = 0.17). Bees colonized with the BeeCom_001 showed a lower SRS compared to bees mono-colonized Bifidobacterium (Dunn’s post hoc test corrected with FDR: BC vs. Bi: p < 0.01). Despite a higher sucrose responsiveness, bees mono-colonized with Bifidobacterium performed poorer in the olfactory conditioning assay compared to bees colonized with the BeeCom_001, confirming differences in learning and memory processes between these gut conditions.

## Discussion

Our study revealed that the core community of gut bacteria common to corbiculate bees facilitates honey bees’ ability to discriminate odours based on their possible association with a sucrose reward. The bacterial community did not affect the perception of that reward as demonstrated by similar sucrose responsiveness between GF bees, mono-colonized bees and bees colonized with the BeeCom_001. Different performances in the olfactory conditioning assay were therefore not explained by differences in hunger or motivation but rather by the modulation of olfactory perception, learning and memory processes by gut bacteria. Interestingly, colonization with individual community members could not reproduce the effect of the community on bees’ cognitive performance. None of these members’ specific abundance was associated with success or failure in the memory test. Furthermore, the total number of bacteria in the gut of bees colonized with the BeeCom_001 did not differ from bees mono-colonized with Bombilactobacillus Firm-4 and Lactobacillus Firm-5 and could therefore not explain the different levels of cognitive performance between these gut conditions. Altogether, it suggests that cognitive improvements were an emergent property of the community.

Defined communities have improved our understanding of inter-species interactions among bacterial members as well as trans-kingdom communication in host-microbe symbioses^21,22^. In *Drosophila melanogaster*, the exchange of metabolites between community members influenced their host’s olfactory preferences and egg-laying behaviour in a different manner compared to individual members^23^. Similarly, we did not see any effect of individual members on bees’ performance suggesting that interaction between members trigger the cognitive improvements. While tryptophan degradation by Lactobacillus Firm-5 was previously shown to increase olfactory memory, such an effect of Firm-5 in absence of tryptophan supplementation was not observed in mono-colonized bees^10^. One hypothesis could be that Firm-5 utilizes the tryptophan produced by other bacterial species in the community. Tryptophan was indeed found more abundant in the gut of bees colonized with a natural community than in GF bees^10,24^. Elevated abundances of the bacterium Lactobacillus Firm-5 within the natural gut community was also shown to promote visual memory performances in bumblebees^25^. However, in that study, glycerophospholipid production was identified as the key metabolic pathway facilitating memory formation. Further investigations will help identify the metabolites resulting from inter-species interaction within the BeeCom_001 that are responsible for the observed cognitive improvements of host bees.

There are multiple ways in which the bacterial products of the BeeCom_001 may promote olfactory learning and memory performance. First, they might participate in brain maturation and cognitive development, as observed in humans and mice^26,27^. Honey bees acquire their gut microbiota during perinatal life, concomitantly to structural and functional brain maturation^28,29^. An optimal capacity to memorize odour—food associations was observed between 5-8 days after emergence, the age at which we tested bees’ performance in our assay^30,31^. Gut colonization by the BeeCom_001 during the first week of adulthood might have facilitated processes involved in brain maturation which resulted in the reported cognitive improvement. Second, and independently of cognitive development, bacterial products might facilitate the neuronal plasticity triggered by the coincidence detection of the CS and US and required for the formation of memories^32,33^. Adult mice exposed to an antibiotic inducing a dysbiosis in the gut showed reduced learning-related structural and functional plasticity of the neurons involved^8^. Further investigations are needed in honey bees to discriminate the effect of the BeeCom_001 on cognitive development from effects on learning processes *per se*.

Finally, gut bacteria may also be susceptible to affect indirectly learning and memory by modulating bees’ appetite for sugar. Previous studies indeed reported an increased sucrose response score in bees colonized with a natural gut microbial community compared to GF bees^24,34^, and olfactory learning performance measured in the conditioning of the PER assay was shown to correlate positively with sucrose response scores^35^. Yet, sucrose responsiveness was not impacted by the gut condition in our study, eliminating the possibility that the observed cognitive improvements in bees colonized with the BeeCom_001 are due to the US perception or motivation for food. The contradiction with previous studies may come from their utilisation of a gut homogenate of hive bees to colonize the gut of the tested bees^24^. This homogenate contains a greater diversity of micro-organisms as well as pollen fermented compounds that are absent from our BeeCom_001 inoculum. The discrepancy could also have arisen from different starvation times (2h in^36^, 18h in ours). Starvation time was indeed shown to have a critical influence on sucrose responsiveness, appetitive learning and memory^37^. Finally, there are methodological discrepancies with the previous study by Zhang et al.^34^: (i) they considered the first presentation of water as a sucrose dilution, which indicates thirst or overall sensitivity to antennal stimulation rather than sucrose responsiveness; (ii) they eliminated bees that did not respond to the highest sucrose concentration, which is a variable that can be affected by the bacterial treatment and is part of the sucrose responsiveness assay; and (iii) they assumed that bees responding to a given concentration of sucrose would also respond to higher concentrations when computing the SRS, which ignores the possibility that some bees responded randomly to the presented stimuli. The method used by Zhang et al.^34^ might have led to an overestimation of the SRS in colonized bees who massively responded to the first presentation of water. An analyses of water responses may have revealed differences in habituation and sensitization to antennal stimulation accounting for the differences in SRS.

Our findings on the positive impact of the defined corbiculate core community on honey bee cognition open new avenues of research on the gut microbiota – brain – axis. The value of the BeeCom_001 as a probiotic should also be tested in bees suffering from dysbiosis induced by environmental stressors^38^ in an effort to help protect threatened bee populations. Finally, the ecological relevance of the BeeCom_001 opens evolutionary questions regarding the selection pressures that apply to corbiculate bees’ cognition and gut bacteria and which have shaped this symbiotic association^39^.

## Materials and Methods

### Generation of gnotobiotic bees

Germ-free honey bees were obtained for 14 experimental replicates each from a different colony of *Apis mellifera carnica* maintained on the campus of the University of Lausanne during the spring and summer seasons of 2021 and 2022. They were generated as described in Kešnerová et al.^17^ and randomly assigned to one of the following treatment groups: germ-free (GF), colonized by the defined corbiculate core community BeeCom_001, mono-colonized by Gilliamella, Snodgrassella, Bombilactobacillus Firm-4, Lactobacillus Firm-5, or Bifidobacterium.

Bacterial strains used for the colonization were obtained from glycerol stocks stored at −80°C. Each bacterial strain was cultured under specific conditions, restreaked once, washed in 1X PBS, diluted to OD_600_ = 1 and stored in 20% glycerol at −80°C until further use. Details on bacterial strains and culturing conditions can be found in the Supplementary Table 1. Colonized bees were obtained by feeding GF bees 5μL of a solution containing the colonization stocks diluted ten times in a 1:1 mixture of 1X PBS and 50% sucrose solution (w/v). The colonization stock used for the GF group consisted in 20% glycerol in 1X PBS. For mono-colonization, strains belonging to the same phylum (e.g. Gilliamella) were mixed in equal proportion before being diluted in the PBS/sucrose solution. The BeeCom_001 contained all eleven strains in equal proportions.

Colonized bees were housed in cages of 10-19 individuals depending on the mortality encountered during the generation of GF bees. For a given replicate, all treatment groups contained the same number of individuals. Bees had unlimited access to 50% sucrose solution (w/v) and to sterilized pollen. Cages were kept at 30°C with 70% humidity for seven days. On the last evening, bees were transferred to clean cages for a 12-hours overnight fasting.

### Sucrose responsiveness assay

Sucrose responsiveness was assessed by recording proboscis extension responses to increasing concentrations of sucrose, as described in Scheiner et al.^35^. On the eighth day post-colonization, bees were anesthetized on ice and harnessed in 3D-printed tubes allowing movements of the antennae and mouthparts only. Harnessed bees were kept resting for 2 h. Bees were presented an ascending concentration series of sucrose (i.e. 0.1%, 0.3%, 1.0%, 3.0%, 10%, 30% w/v) by touching their antennae with the corresponding solution. Each stimulation with sucrose was preceded by a stimulation with water. The interval between two stimulations of the antennae was 3 min. The presence or absence of PER to each stimulation was recorded. A subset of bees responding to the highest sucrose concentration, but not to all water stimulations, was randomly selected for the olfactory conditioning assay.

### Olfactory conditioning

The protocol for the olfactory conditioning of the PER was adapted from Matsumoto et al.^19^ and started after 1 h of rest. The experimenter was blinded with respect to the treatment group. Bees were trained to discriminate an odour A associated with a sucrose reward from an unrewarded odour B. The odours heptanal and 1-nonanol (Sigma-Aldrich) were used alternately as A or B between each replicate. They were chosen based on their level of perceptual similarity^40^. The conditioning assay included in five trials of 40 s for each odour, presented in a pseudo-randomised order. For each trial, bees were placed on the conditioning set-up, in front of a syringe containing a filter paper soaked with 5 μL of pure odorant. After 12 seconds of familiarization with the context, a constant airflow was sent through the syringe, thereby delivering the odour to the bee for 4 s. A toothpick soaked in 50% sucrose solution (w/v) was presented to the bees’ antennae 3 s after the onset of odour A delivery. Bees extending their proboscis were then allowed to drink the sucrose solution for 3 s. The intertrial interval was 10 minutes. The presence or absence of PER during the odour presentations was noted as 1 or 0, respectively. Bees responding to the first presentation of the odour A and bees showing no PER to the sucrose stimulations were discarded from the analysis (< 5% of the bees). Short-term memory was assessed by presenting the odours A and B in a randomized order 15 min after the end of the conditioning experiment.

The guts of the tested bees were collected immediately after the memory test and stored at −80°C for further analyses. When possible, one gut per condition and experimental replicate was processed for qPCR analyses.

### DNA extraction from honeybee gut tissue

A total of 80 guts from 13 experimental replicates was used to assess the gut microbiota composition. The protocol for DNA extractions was adapted from Kešnerová et al.^17^. Samples were homogenized with 750μL of deionized water, zirconia beads (0.1 mm dia. Zirconia/Silica beads; Carl Roth) and glass beads in a Fast-Prep24 5G homogenizer (MP Biomedicals) at 6 m/s for 45 s. Half of the gut homogenate was removed and stored at −80°C as a backup. Samples (still containing the beads) were homogenized again with 375 μL of CTAB lysis buffer (0.2M Tris-HCl, pH 8; 2.8 M NaCl; 0.04 M EDTA, pH 8; 4% CTAB, w/v, dissolved at 56°C; 2 μL β-mercaptoethanol; 20 μL proteinase K [20mg/mL]) and incubated at 56°C for 1 h. Homogenates were mixed with 750 μL phenol:chloroform:isoamyl alcohol (Fischer Bioreagents, pH 8), centrifuged at 16,000 x g for 10 mins at room temperature. The upper aqueous phase (500 μl) was transferred to a tube containing the same volume of chloroform, mixed and centrifuged again at 16,000 x g for 10 mins at room temperature. The upper aqueous phase (500 μl) was mixed with 900 μl of pre-cooled 100% ethanol and incubated overnight at −20°C for precipitation of nucleic acids. After a centrifugation at 16000 x g at 4°C for 30 min, the pellets were washed with 900 μl of 70% ethanol, left to dry at room temperature and resuspended in 50 μl of nuclease-free water (Invitrogen) by shaking in a thermomixer (64°C, 400 rpm, 10 min). The purification of nucleic acids was performed using CleanNGS magnetic beads (CNGS-0005) and the Opentron OT-2 pipetting robot. Purified DNA extracts were stored at −20°C until further use.

### Quantification of bacterial loads in the gut

All qPCR reactions were carried out in a 96-well plate on a QuantStudio™ 5 (Applied Biosystems). The thermal cycling conditions were as follows: denaturation stage at 50°C for 2 min followed by 95°C for 2 min, 40 amplification cycles at 95°C for 15 s, and 60°C for 1 min. Each reaction was performed in triplicate in a total volume of 10 μL (0.2 μM of each forward and reverse primer; 1x SYBR^®^ Select Master Mix, Applied Biosystems; 1 μL DNA). Each DNA sample was screened with two different sets of primers targeting either the actin gene of *A. mellifera*, or the universal 16S rRNA region. Samples from GF bees and bees colonized with the BeeCom_001 were screened with the five sets of species-specific primers. Samples from mono-colonized bees were screened with the set of species-specific primer corresponding to the phylotype present in their inoculum. Information about primers and absolute quantification method can be found in Kešnerová et al.^17^.

### Statistical analyses

All statistical analyses were performed with R Studio 2022.02.0^41^. The presence or absence of PER to the CS during the olfactory conditioning and memory test was recorded as a binomial variable. Different generalized linear mixed models with a binomial error structure — logit-link function — (glmer function of R package lme4) were used to assess the predictive value of the trials and gut condition on responses to the CS (conditioning assay) and to the stimuli (sucrose responsiveness assay). The GF gut condition was used as the reference level for all models. Individual identity was set as a random factor. Tukey *post hoc* analyses were run whenever a statistical difference was detected in the models. Kruskal-Wallis test, followed by pairwise Wilcoxon Rank Sum tests corrected with the false-discovery rate method were used to compare mortality, sucrose consumption, carcass weight and gustatory response scores (SRS and WRS) between the different gut conditions.

## Acknowledgements

We thank Armindo Teixeira for building the odour delivery system, Paul Marchal for the design of the 3D-printed harnessing tubes and Matthieu Vollmer for helping with generating gnotobiotic bees. This work was funded by the University of Lausanne, the Marie Skłodowska-Curie fellowship HarmHoney (grant no. 892574, awarded to A.C.), the ERC Starting Grant MicroBeeOme (grant no. 714804, awarded to P.E.), and a Swiss National Science Foundation project grant (grant no. 179487, awarded to P.E.), and the NCCR Microbiomes, a National Centre of Competence in Research, funded by the Swiss National Science Foundation (grant no. 180575).

## Notes

### Competing Interest Statement

The authors have declared no competing interest.

