## Supplementary Information for "A defined community of core gut microbiota members promotes cognitive performance in honey bees"

**Supplementary Table 1. Bacterial strains used in this study and their culturing conditions.** The proportion of each strain relative to other strains present in the inoculum of the different gut treatments is provided under brackets. The defined community BeeCom_001 (BC) contained all 11 strains.

| **Bacterial species** | **Strain name** | **Gut treatments** | **Culturing condition** |
|---|---|---|---|
| *Gilliamella apicola* | ESL0178 | BC (1:11) and Gi (1:3) | BHIA, 35°C, 5% $CO_2$ |
| *Gilliamella apis* | ESL0169 | BC (1:11) and Gi (1:3) | BHIA, 35°C, 5% $CO_2$ |
| *Gilliamella sp.* | ESL0177 | BC (1:11) and Gi (1:3) | BHIA, 35°C, 5% $CO_2$ |
| *Snodgrassella alvi* | ESL0145 (wkB2) | BC (1:11) and Sn (1:1) | TSA, 35°C, 5% CO2 |
| *Lactobacillus mellis* (Firm-4) | ESL0094 (Hon2N) | BC (1:11) and F4 (1:1) | MRSA, 37°C, anaerobic |
| *Lactobacillus apis* (Firm-5) | ESL0185 (Hma11) | BC (1:11) and F5 (1:4) | MRSA, 37°C, anaerobic |
| *Lactobacillus helsingborgensis* (Firm-5) | ESL0183 (Bma5) | BC (1:11) and F5 (1:4) | MRSA, 37°C, anaerobic |
| *Lactobacillus melliventris* (Firm-5) | ESL0184 (Hma8) | BC (1:11) and F5 (1:4) | MRSA, 37°C, anaerobic |
| *Lactobacillus kullabergensis* (Firm-5) | ESL0186 (Biut2) | BC (1:11) and F5 (1:4) | MRSA, 37°C, anaerobic |
| *Bifidobacterium asteroides* | ESL0170 | BC (1:11) and Bi (1:2) | MRSA, 37°C, anaerobic |
| *Bifidobacterium asteroides* | ESL0197 | BC (1:11) and Bi (1:2) | MRSA, 37°C, anaerobic |

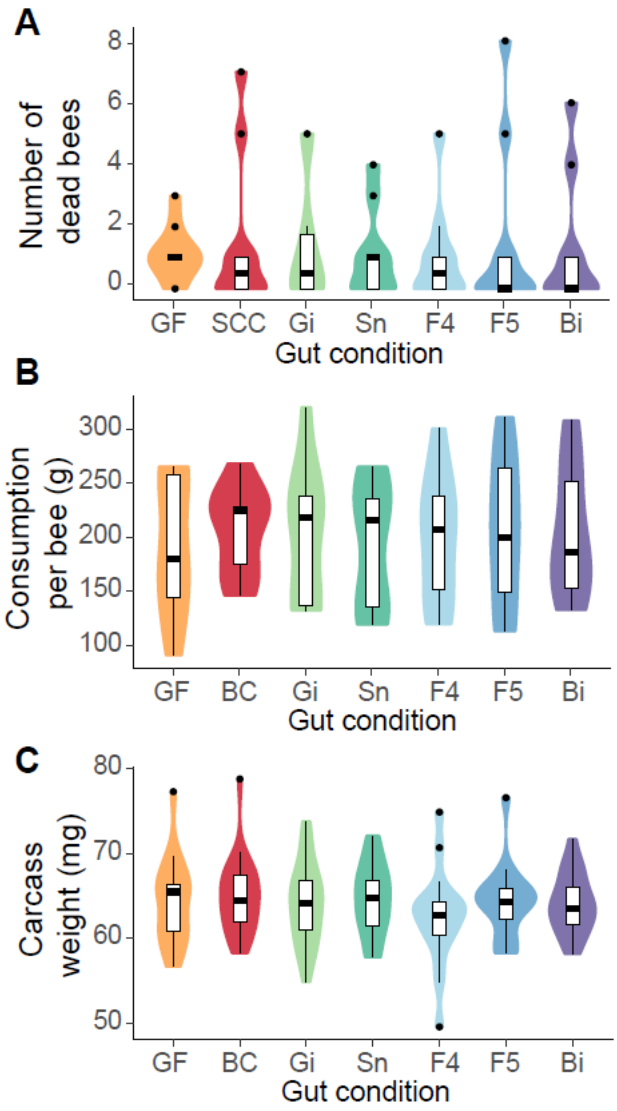


**Supplementary Figure S1. Impact of the gut condition on bees' physiology. (A)** The number of dead bees and **(B)** the sucrose consumption in the rearing cages was recorded for 7 days (n = 10 cages per gut condition). **(C)** The carcass weight of bees was measured following gut dissections. The gut conditions were germ-free (GF, n = 21), colonized with the synthetic core community BeeCom_001 (BC, n = 29), and mono-colonized with Gilliamella (Gi, n = 23), Snodgrassella (Sn, n = 21), Bombilactobacillus Firm-4 (F4, n = 19), Lactobacillus Firm-5 (F5, n = 20), or Bifidobacterium (Bi, n = 22).

Kruskall-Wallis tests were non-significant (Mortality: $\chi^2 = 1.43$, df = 6, p = 0.96; Consumption: $\chi^2 = 0.6$, df = 6, p = 1; Carcass weight: $\chi^2 = 3.37$, df = 6, p = 0.76).

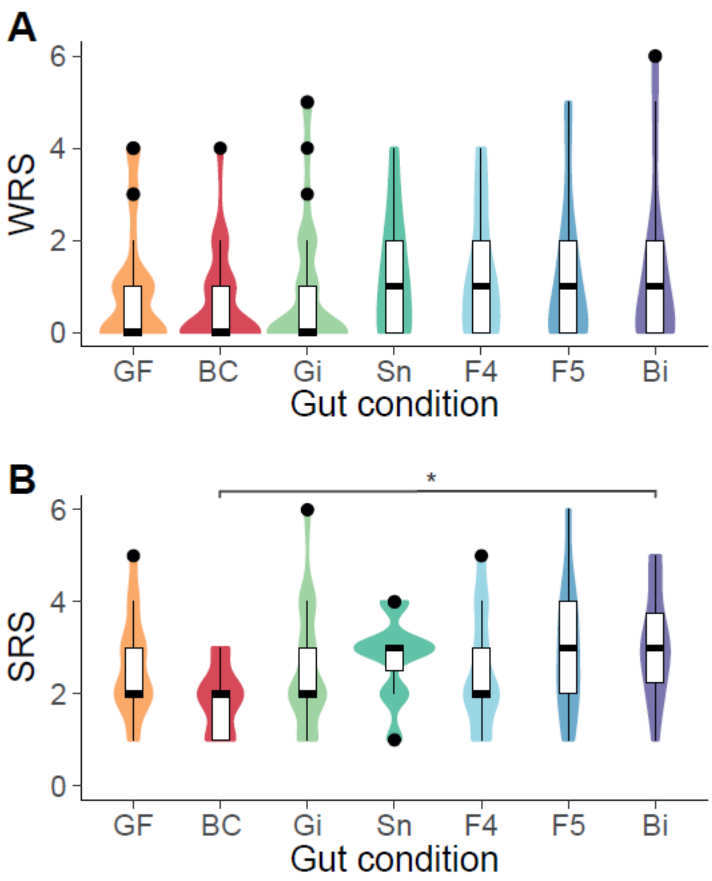


**Supplementary Figure S2. Sucrose and water response scores of gnotobiotic bees that were selected for the olfactory conditioning assay.** The gut conditions were germ-free (GF, $n_A$ = 35, $n_B$ = 18), colonized with the synthetic core community BeeCom_001 (BC, $n_A$ = 35, $n_B$ = 22), and mono-colonized with Gilliamella (Gi, $n_A$ = 39, $n_B$ = 24), Snodgrassella (Sn; $n_A$ = 31, $n_B$ = 11), Bombilactobacillus Firm-4 (F4; $n_A$ = 37, $n_B$ = 14), Lactobacillus Firm-5 (F5; $n_A$ = 33, $n_B$ = 16), or Bifidobacterium (Bi; $n_A$ = 37, $n_B$ = 18). (*) $p < 0.05$, Dunn's post hoc test corrected with the false-discovery rate method.
